# For running or burying—The toe fringe of *Phrynocephalus mystaceus* is important

**DOI:** 10.1101/2019.12.26.889105

**Authors:** Peng Zheng, Tao Liang, Jing An, Lei Shi

## Abstract

Toe fringe is the most typical morphological feature of lizards adapted to sandy environments, and toe fringe is suitable for testing the adaptive convergence suggested by form-environment correlations. *Phrynocephalus mystaceus* mainly lives in dune habitats, has a developed bilateral toe fringe, and exhibits fast sand-diving behavior for predator avoidance. We tested the effects of resecting the medial and bilateral toe fringes on the locomotor performance and sand-diving performance of *P. mystaceus.* The results showed that the individuals that had their medial toe fringe removed exhibited significantly faster sprint speeds than the unresected and all resected individuals (*P* <0.05). The results of stepwise regression analysis show that the relative area of the toe fringe is an important morphological feature that affects locomotor performance. The sand-burial performance scores of the unresected individuals were significantly greater than those of the resected individuals (*P* <0.05). The results of stepwise regression analysis show that the relative area of toe fringe and the axilla-groin length are the main morphological features that affect the sand-diving performance of *P. mystaceus*. After the bilateral toe fringe was removed, a significant negative correlation between locomotor and sand-diving performance was observed (*P* <0.05). Taken together, these results provide experimental evidence that toe fringe is positively associated with the locomotor and sand-diving performance of *P. mystaceus*.

---

Locomotion is a fundamental component of prey capture (Higham, 2007). An animal’s escape behavior should reflect both the cost of interrupting current activities to respond to predators and the relative risk of predation (Ydenberg and Dill, 1986; Cooper and Frederick, 2007). Some lizards escape predators by rapidly burying themselves in sand (Darwin 1962; Arnold 1995; Attum et al., 2007; Kacoliris et al., 2010).

Morphological differences can explain the diversity of behavior in many species (Arnold, 1983). Similar morphological characteristics for adaptation to the desert environment have evolved in different groups of desert lizards. One of the most common morphological characteristics that has evolved is toe fringe. “Lizard toe fringes are composed of laterally projecting elongated scales and have arisen independently at least 26 times in seven families of lizards” (Luke, 1986). According to the report, lizards with toe fringe run faster than those without toe fringe on sand surfaces; nevertheless, lizards with toe fringe exhibit slow speeds on rubber surfaces compared with individuals without fringe (Carothers, 1986). Except for locomotion abilities, toe fringe can also explain sand-diving, a behavior used for submersion into granular substrates (Stebbins, 1944). However, it is not clear exactly what role fringe plays in the locomotor behavior of lizards. *Uma scoparia* have well-developed fringes, which are thought to improve sprinting performance over fine sand (Irschick and Jayne, 1998; Korff and McHenry, 2011). Studies have shown that the locomotor performance of lizards on sand significantly decreased after the fringes were removed. In particular, the locomotor performance of lizards was significantly decreased on the uphill sloped surfaces (Carothers, 1986). Other studies have compared *Uma scoparia*, which has fringe, to *Callisaurus draconoides*, which does not have fringe. These studies found no significant differences in the sprint speeds on the different substrates (Bergmann and Irschick, 2010; Korff and McHenry, 2011; Li et al., 2012). On the other hand, with the decrease in substrate resistance, the performance of *C. draconoides* did not decrease significantly, but that of *U. scoparia* did (Qian et al., 2015). These studies indicate that fringes may perform a variety of functions.

For instance, *U. scoparia* uses its fringe for fast sand-diving behavior to escape predators and extreme heat (Arnold, 1995; Attum et al, 2007). In addition, *C. draconoides* and *U. scoparia* belong to different genera and show great differences in the relative limb proportions and potential behavioral, ecological, and physiological aspects. This uncontrolled variation complicates the examination of interspecific performance (Carothers, 1986).

*Phrynocephalus mystaceus* is the largest species of *Phrynocephalus*, which is a genus of toad-headed agama lizards (Zhao et al., 1999; Solovyeva et al., 2018). This species is also a typical desert lizard species from central Asia to northwest China, and it mainly lives in dune habitats and has a well-developed bilateral triangle toe fringe (Luke, 1986). *P. mystaceus* can run quickly over fine sand substrates and exhibits fast sand-diving behavior when avoiding predators (Arnold, 1995). There is a lack of research on the morphology and locomotor performance of *P. mystaceus*. This species is listed as endangered on the Red List of China’s Vertebrates (Jiang, 2016). Therefore, studying the locomotor behavior of this species may help us better protect *P. mystaceus.* While there have been reports of studies that have removed the toe fringes of sand-dwelling lizards to test locomotor performance (Carothers, 1986), there are few reports of studies that have removed the toe fringes of sand-dwelling lizards to test their sand-diving performance. In this study, we measured several morphological traits and analyzed the locomotor and sand-diving performance of *P. mystaceus* on sand substrates. In particular, we adopted a control test that consisted of removing the toe fringes to verify the following scientific hypotheses: (1) The presence or absence of toe fringes on *P. mystaceus* will affect its locomotor performance over sand substrates. (2) The presence or absence of toe fringes on *P. mystaceus* will affect its sand-diving performance on sand substrates. (3) The toe fringes of *P. mystaceus* influence the locomotion and sand-diving performance, and there is a trade-off between the two performances.

## Materials and methods

In July 2018, we collected *P. mystaceus* individuals by hand from the Tukai Desert, Huocheng County, Yili Region, Xinjiang. The selected individuals in good condition were taken back to the Zoology Laboratory of Xinjiang Agricultural University. We measured the snout-vent length (*X*_1_: SVL), head length (*X*_2_: HL), head width (*X*_3_: HW), head depth (*X*_4_: HD), mouth breadth (*X*_5_: MB), axilla-groin length (*X*_6_: AG), abdominal width (*X*_7_: AW), tail base width (*X*_8_: TBW), fore limb length (*X*_9_: FLL), hind limb length (*X*_10_: HLL) and tail length (*X*_11_: TL) (Zhao, 1999). All measurements were accurate to within 0.1 mm. The toe fringes of the lizards were quantified according to the following characteristic traits: individuals’ total fringe number divided by snout-vent length (*X*_12_:TFN), individuals’ total fringe max length divided by snout-vent length (*X*_13_:TFL), and individuals’ total fringe area divided by snout-vent length (*X*_14_:TFA). We measured the toe fringe characteristics with a Canon digital camera and then analyzed data with image-pro Premier 6.0 software. *P. mystaceus* individuals were kept in tanks for lizards. The tanks were covered with 5 cm of fine sand collected from the original habitat of *P. mystaceus*, with a 60-w bulb suspended at one end as a heat source for thermoregulation. Plenty of *Tenebrio molitor* larvae and water supplemented with calcium and vitamins were provided to ensure that the animal received a full complement of nutrients. The animals were kept temporarily for 1 week before beginning the exercise test, and all tests were completed within 2 weeks. All animals were released to the original capture site after the test.

Locomotion performance was measured on a 1.4 m horizontal track, and the racetrack was covered with sand substrate from the original habitat. Before testing the locomotion performance, we conducted a preliminary test to determine the optimal temperature for the activity of *P. mystaceus*. A temperature gradient was designed from 0 □ to 50 □, and it was found that the optimal temperature for the activities of the great oared lizard was approximately 34 □; thus, before the locomotion performance test, all individuals were preconditioned for 1 h at (34.0±0.5) □. Then, we moved the animal into the end of the track, and a brush was used to push it to sprint. A digital camera was used to record the lizard’s movements on the track. Through video playback, the 1.4 m track was divided into seven segments. In addition, the frames of each segment were counted, the motion time in the video was analyzed by Adobe premiere CS6 software, and the motion velocity was calculated (Higham et al., 2010). The speed was graded by using an arithmetic sequence (to be divided by ten classes). The entire exercise test was divided into three repeats: no cut fringes, single cut (removal of the medial fringes) and double cut (removal of bilateral fringes).

The sand-diving performance was measured in a tank covered with 10 cm of fine sand from the original habitat. We used a brush to stimulate the tail, which caused the individuals to dive into the sand. The entire process was recorded with a digital camera, and sand-diving behavior was recorded through video playback. The whole sand-burying behavior was also divided into three repetitions: no cut fringes, single cut (removal of the medial fringes) and double cut (removal of bilateral fringes). The sand-burying time was graded by an arithmetic sequence.

The scores of sand-burying behavior were as follows: sand-burying ability score, sand-burying time score and comprehensive score. Among them, the sand-burying ability score was based on the sand-burying state of *P. mystaceus*: fully buried: 5 points, tail not buried: 4 points, head not buried: 3 points, most of the body not buried: 2 points, not buried in the sand: 1 point. The sand-burying time score was used to sort the burying time from small to large according to the arithmetic sequence and was divided into 5 grades, with the shortest burial time assigned a value of 5 points, and the points were successively divided into 4 points, 3 points, 2 points and 1 point according to the sorting method of the arithmetic sequence. Then, the comprehensive score was the sum of the two score types. Individuals with the best and fastest burial performance had the highest the overall scores.

### Statistical analyses

The data were tested by Kolmogorov–Smirnov tests for detecting normality. We log-transformed the variables to minimize the heterogeneity, where necessary (King, 2000). We used analysis of covariance (ANCOVA) to examine the differences in locomotion performance and sand-diving performance of the different states on the sand substrates with paired sample t tests (multiple comparisons). We used stepwise regression analysis to screen the morphological characteristics that could be used to determine the movement ability over sand substrate and the morphological characteristics that could be used to determine the buried sand score for sand-diving behavior. We used Fisher’s exact test to examine the differences in the probability of sand diving under different states. Spearman correlation was used to analyze the correlation between the velocity score and the comprehensive score. All analyses were conducted using R v. 3.5.1 (R Core Team, 2017).

## Results

### Locomotion performance

In the running trials, there were significant differences in the maximum sprint speeds on the sand substrate under the different conditions (repeated measures ANOVA, *F*_2, 8_=4.524, *P*=0.02). In addition, the results of multiple comparisons (the paired sample t test) showed that the maximum sprint speed after removing the medial fringes was significantly higher than that before removing the medial fringes (*P*<0.05, Fig. 1). In addition, there were no significant differences in the sprint speeds under the other states (*P*>0.05, Fig. 1). Multiple stepwise regression analysis showed that under the no cut state, TFA (*X*_14_) was the major determinant of locomotion performance. Under the single cut state, TFN (*X*_12_) and TFA (*X*_14_) were the major determinants of locomotion performance. However, under the double cut state, AG (*X*_6_) and TBW (*X*_8_) were the major determinants of locomotion performance (Table 1).

**Table 1.**
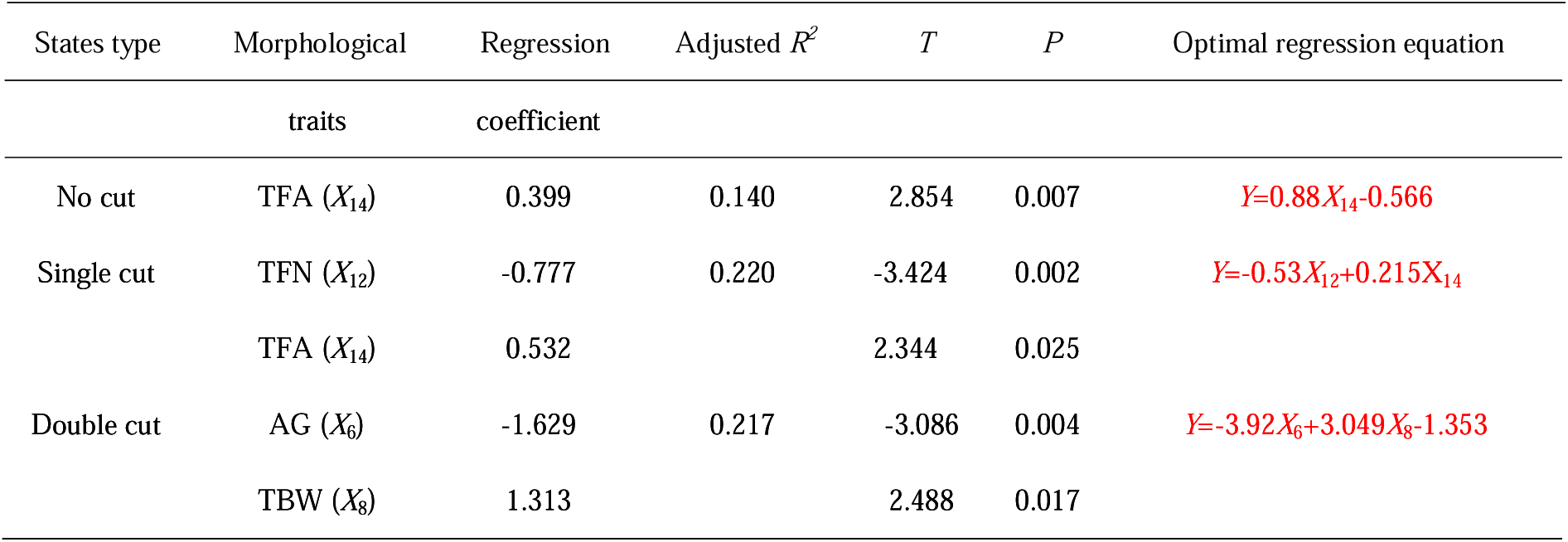
Stepwise regression analysis results of the morphological traits affecting the maximum sprint speed under different states

**Fig. 1.**
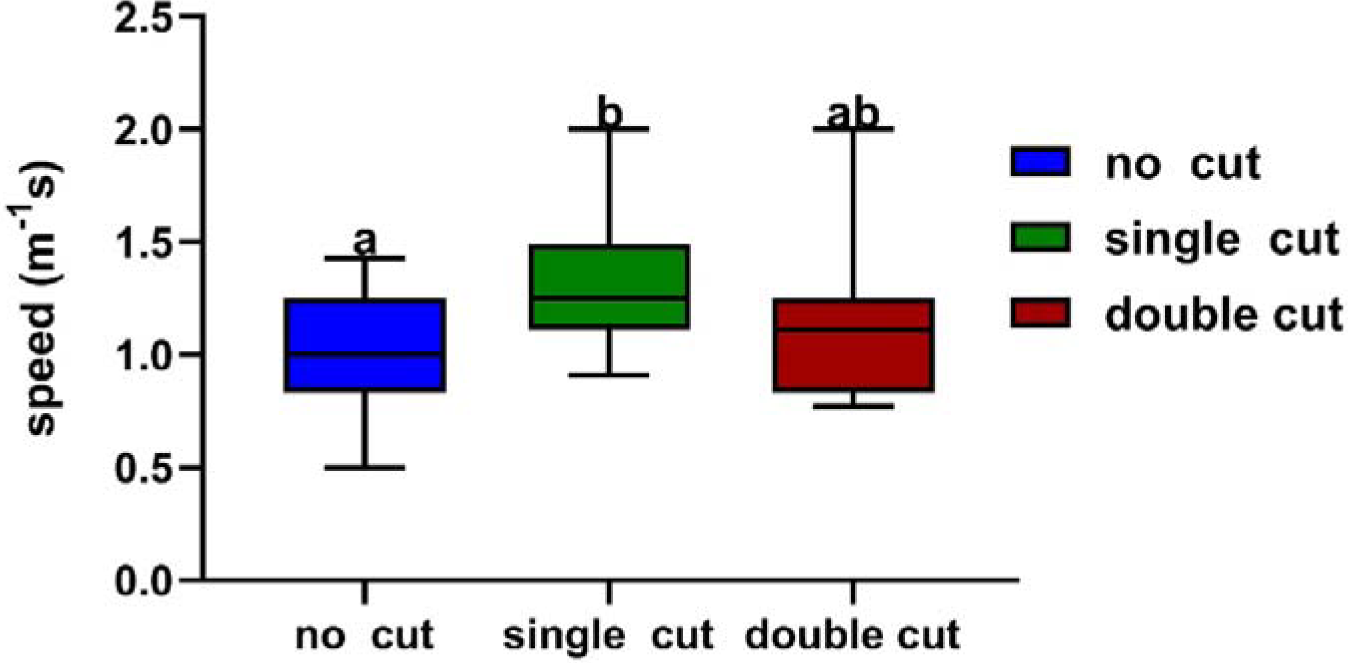
The maximum sprint speed over sand substrate of *Phrynocephalus mystaceus* under different states. Note:Different letters indicate significant differences at the *P* < 0.05 level

### Sand-diving performance

The results of the univariate repeated measures ANOVA by sphericity test show that under different states, there were no significant differences in the sand-burying ability scores on the sand substrate (repeated measures ANOVA, *F*_2, 8_=2.057, *P*=0.171). However, the sand-burying ability scores after removing bilateral fringes were significantly lower than those before cutting (*P*<0.05, Fig. 2). In addition, there were no significant differences in the sand-burying ability scores under the other states (*P*>0.05, Fig. 2). There were no significant differences in the sand-burying time scores on the sand substrate (repeated measures ANOVA, *F*_2, 8_=3.019, *P*=0.077). However, the sand-burying time scores after removing bilateral fringes were significantly lower than those before cutting (*P*<0.05, Fig. 3). In addition, there were no significant differences in the sand-burying time scores under the other states (*P*>0.05, Fig. 3). In terms of the probability of sand diving, there were no significant differences under the different states (*P*>0.05, Table 2).

**Table 2.** Descriptive statistics of the ability score frequencies for sand diving by *Phrynocephalus mystaceus* under different states

| States | Sand diving (Total) | No sand diving (Total) | Grouping | $X^2$ | $P$ |
| --- | --- | --- | --- | --- | --- |
| No cut | 9 (9) | 0 (9) | No cut vs Single cut | 2.25 | 0.235 |
| Proportion | 100 | 0 |  |  |  |
| Single cut | 7 (9) | 2 (9) | No cut vs Double cut | 2.25 | 0.235 |
| Proportion | 77.8 | 22.2 |  |  |  |
| Double cut | 7 (9) | 2 (9) | Single cut vs Double cut | 0 | 0.712 |
| Proportion | 77.8 | 22.2 |  |  |  |

**Fig. 2.**
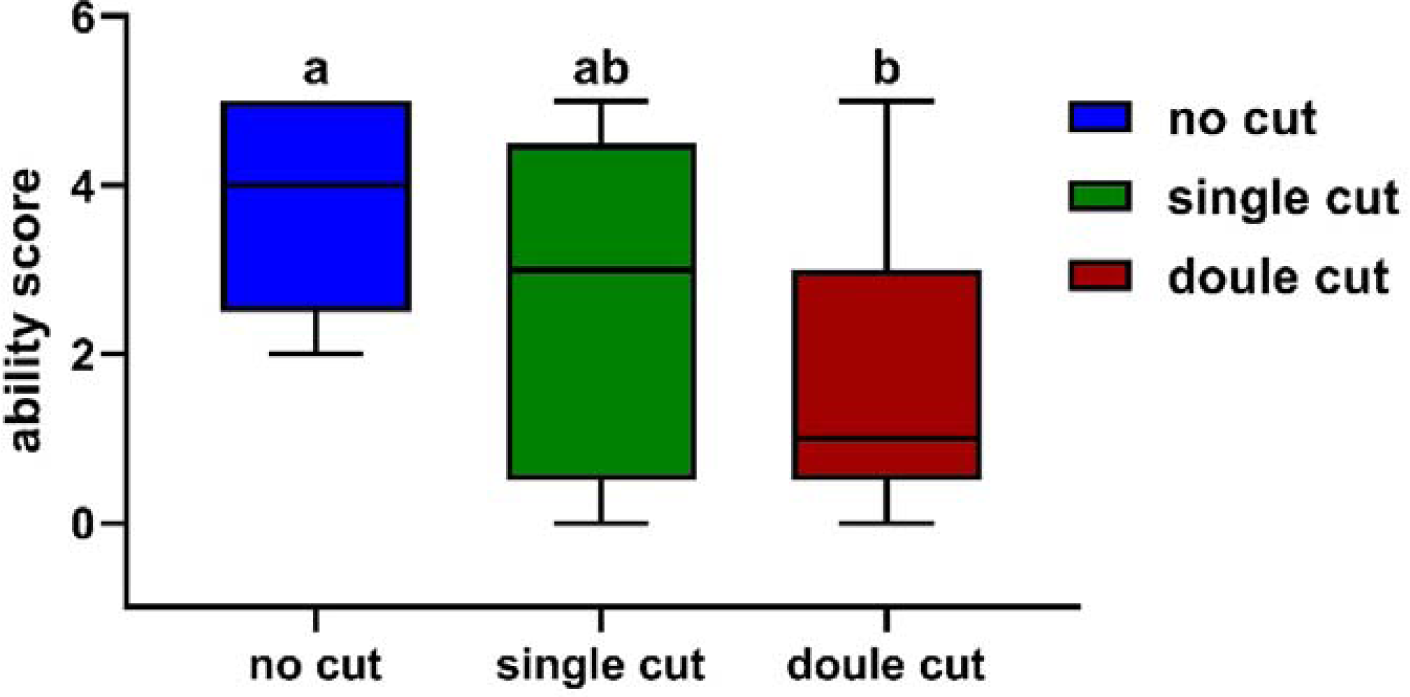
Ability scores for sand diving by *Phrynocephalus mystaceus* under different states. Note:Different letters indicate significant differences at the *P* < 0.05 level

**Fig. 3.**
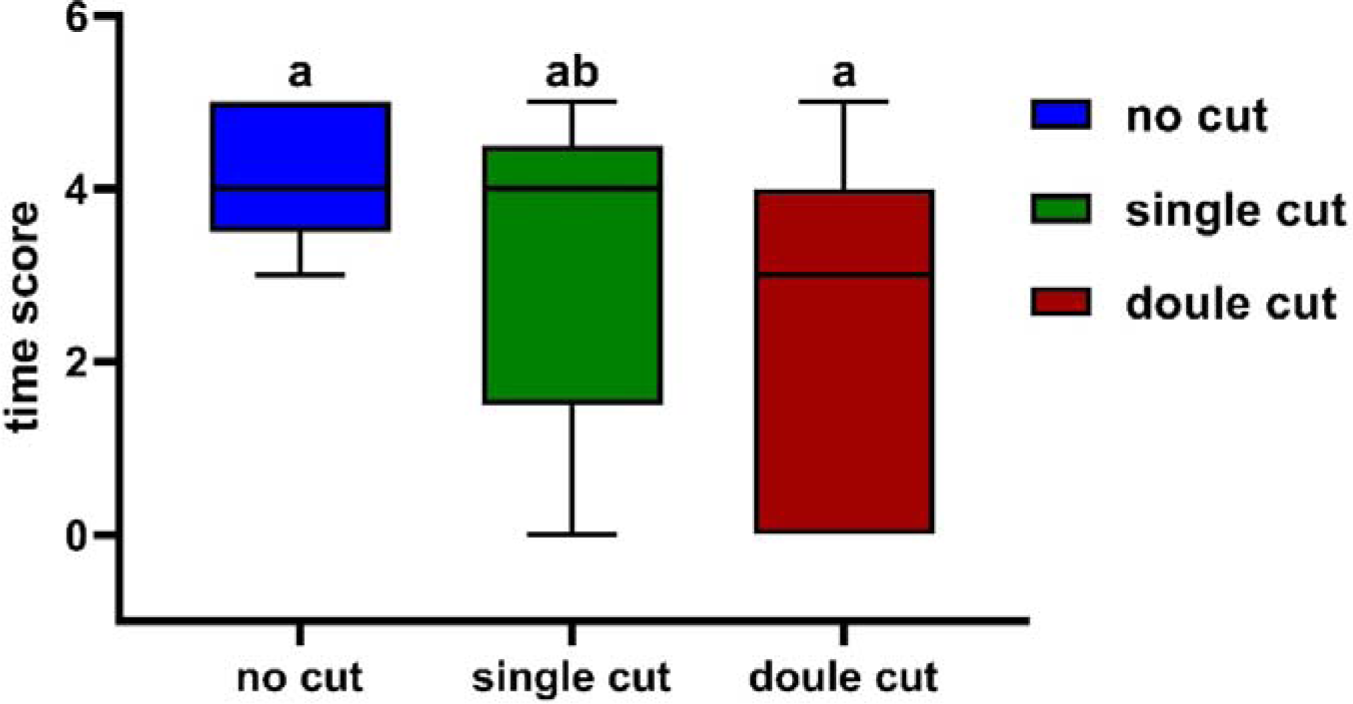
Time scores for sand diving of *Phrynocephalus mystaceus* under different states. Note:Different letters indicate significant differences at the *P* < 0.05 level

In terms of the score type, TFA (*X*_14_) was the major determinant of the sand-burying ability score, and AW (*X*_7_) was the major determinant of the sand-burying ability time. In terms of the type of composite score, AG (*X*_6_) and TFA (*X*_14_) were the key factors (Table 3).

**Table 3.** Stepwise regression analysis results of the morphological traits affecting the different score types

| Score types | Morphological traits | Regression coefficient | Adjusted $R^2$ | $T$ | $P$ | Optimal regression equation |
| --- | --- | --- | --- | --- | --- | --- |
| Ability | TFA ($X_{14}$) | 0.534 | 0.251 | 2.892 | 0.009 | $Y=0.441X_{14}+2.183$ |
| Time | AW ($X_7$) | -0.425 | 0.141 | -2.150 | 0.043 | $Y=-0.189X_7+4.677$ |
| Comprehensive | AG ($X_6$) | 0.623 | 0.387 | 3.602 | 0.002 | $Y=-0.675X_6+0.885X_{14}+6.949$ |
| | TFA ($X_{14}$) | -0.447 | | -2.581 | 0.018 | |

In the case of no cutting, no significant correlations were observed between the velocity score and the comprehensive score (r^2^=0.03, *P*=0.943, Fig. 4A). After removing the medial fringes, there was a negative correlation between the velocity score and the comprehensive score, but the correlation was not significant (r^2^=-0.509, *P*=0.197, Fig. 4B). However, in the case of double cutting, there was a significant negative correlation between the velocity score and the comprehensive score (r^2^=-0.853, *P*=0.015, Fig. 4C).

**Fig. 4.**
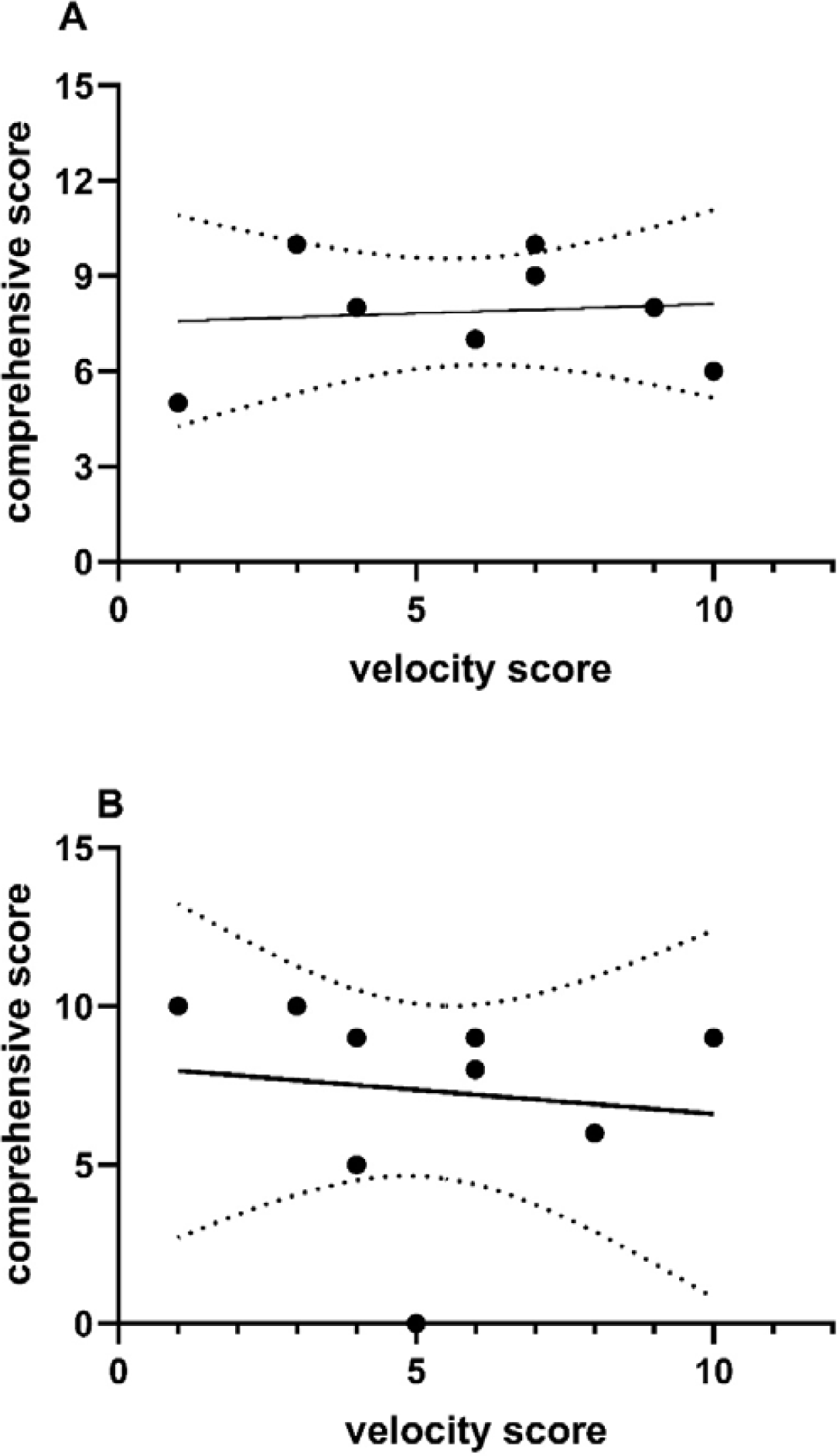
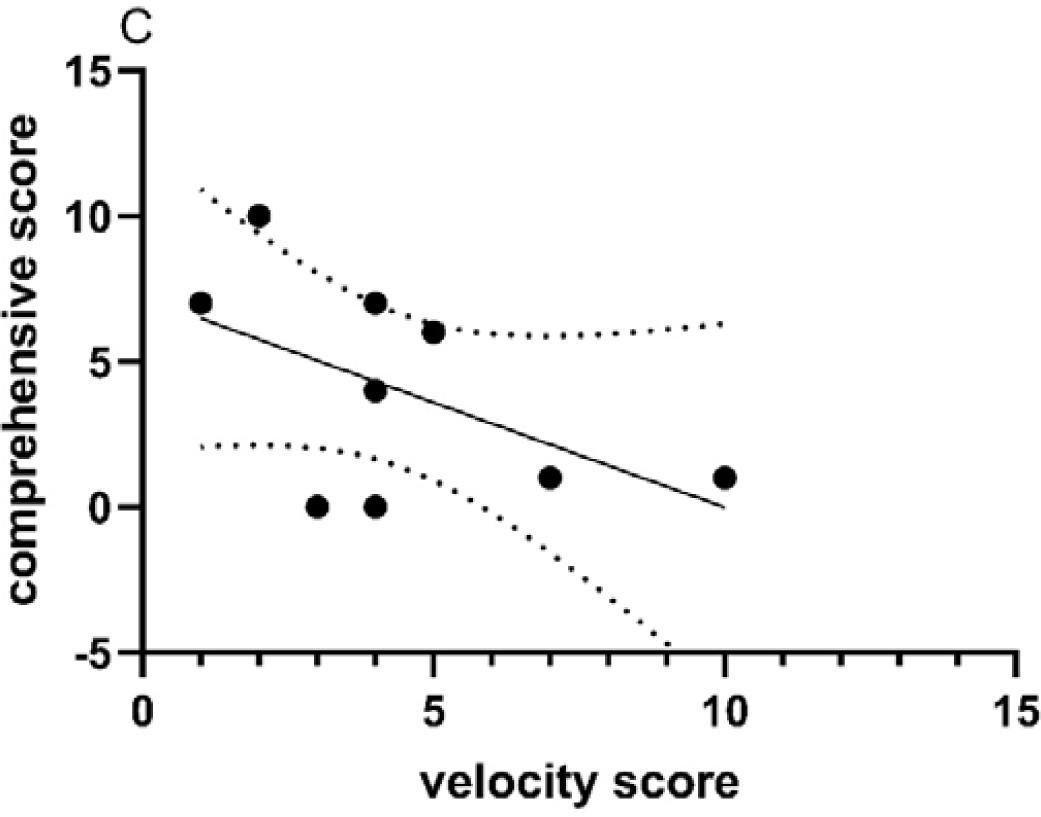
The correlation between the speed score and the comprehensive **sand-diving** score for ***Phrynocephalus mystaceus*** <u>under three states</u> Note: A no cut; B: single cut; C: double cut. curve indicate means with 95% confidence intervals

## Discussion

The morphological characteristics of animals have evolved through natural selection to maximize sprint speed (Irschick and Garland, 2001; Van Damme and Vanhooydonck, 2001). For example, the sprint speed of a lizard is related to body mass and tail size (Ballinger et al., 1979; Downes & Shine, 2001; Du et al., 2005; Johnson et al., 1993; Punzo, 1982). Hind limb length and toe length are also thought to be indispensable factors for a lizard’s sprint speed (Borges-Landáez and Shine, 2003; Vanhooydonck et al., 2002). Natural selection acts on individual variations in locomotor performance in a given environment, thereby altering the trajectory of evolution in the variety of underlying traits governing locomotion (Arnold, 1983; Darwin, 1859; Losos, 2010; Ricklefs & Miles, 1994). In some cases, novel morphological structures have evolved that increase the performance of ecologically relevant tasks (Dornburg et al., 2011; Vermeij, 2006). For example, *R. afer* deploys adhesive toe pads to increase its speed on a level surface (Collins and Higham, 2017). Our results reveal that when *P. mystaceus* is in the no cut or single cut state, the locomotor performance of *P. mystaceus* is related to TFA and TFN (Table 1). In particular, the locomotor performance of lizards on sand significantly increased after the medial fringes had been removed (Fig. 1). However, the locomotor performance of lizards on sand showed a downward trend after the bilateral fringes had been removed (Fig. 1). In addition, the locomotor performance of lizards on the sand did not significantly differ between the no cut and double cut states (Fig. 1). Further explanation is needed for why the outside fringes are one of the main factors that affect the locomotor performance of *P. mystaceus*.

Fringes are often thought to be an adaptation of lizards to allow for sand diving. In *Uma* species, the fringes affect not only the movement of the sand but also the sand-diving ability (Stebbins, 1944). Sand-diving behavior is not only an antipredation behavior (Evans et al., 2017) but also an adaptive behavior that is used to avoid environments that cause water loss and overheating (Halloy et al., 1998; Arnold, 1995). Our results show that under the condition in which sand diving was likely to occur, there were no significant differences under the different states (Table 2). The sand-diving performance when no fringes were cut was significantly higher than when the bilateral fringes had been removed. However, the performance of *P. mystaceus* under the single cut condition was in the intermediate state. This finding further proves that the fringes significantly affected the sand-diving performance. In terms of fringe characteristics, the relative TFA was selected by the regression model of the sand-diving ability score and the comprehensive score for *P. mystaceus*, and these scores were significantly positively correlated (Table 3). This result demonstrates that TFA promotes the sand-diving performance of *P. mystaceus* in terms both scoring types. This is the first time that a control test has been used to confirm that fringes affect sand-diving performance. In addition, there was a significant negative correlation between the sand-diving time score and AW, showing that individuals with larger AW tend to spend more time burying into sand. In some cases, *Phrynosoma* rarely use sprinting as an antipredation strategy, but static antipredation mechanisms such as shape (flat body) and color (body color changes with background color) changes are employed(Stankowich et al., 2016).

At this point, sprint speed is not the deciding factor. The flat body shape of *Phrynocephalus* species is an important antipredation adaptation characteristic, and it is beneficial the hiding ability of the species. However, a larger AW can affect the sprint speed, which may be the result of a trade-off between hiding and escape strategies (zheng et al., 2019). The results of our study on *P. mystaceus* suggest that AW has a negative effect on sand-diving performance and may reflect a trade-off between concealment and sand-diving strategies.

When multiple phenotypic traits perform multiple functions, phenomena such as functional redundancy, tradeoffs, and promotion are ubiquitous (Edwards et al., 2016; Bergmann et al. 2017). Sand diving is a comprehensive antipredation behavior that is related to the escape distance of *P. mystaceus* (Evans, 2017). When confronted with danger in the wild, the lizards that exhibit sand-burying behavior tend to prefer smooth surfaces and soft sand, which may be because it saves energy (Arnold, 1990). However, the antipredator behavior of running is much simpler than sand diving (Arnold, 1990). The trade-off between running and sand diving may be related to the predation pressure. The antipredation strategy is related to the fringe state. When the fringes were not resected, the correlation between the comprehensive score and the velocity score was not obvious. The decision to run or bury in the sand in the face of danger may be related to the flight distance and habitat of *P. mystaceus*. On smooth, soft sand, lizards might be more inclined to bury in the sand. Conversely, the lizards might be inclined to run under different conditions (Evans, 2017). However, after the bilateral fringes were removed, there was a significant negative correlation between running and sand diving, and the sand-diving performance of *P. mystaceus* decreased significantly compared with the locomotor performance. Our results indicate that after removal of the bilateral fringes, *P. mystaceus* preferred to run rather than dive into the sand. It is suggested that the toe fringes of *P. mystaceus* may be important for sand-burying behavior.

Wind-blown debris is the nutritional basis of sand slipface communities (Robinson and Barrows. 2013). Male *P. mystaceus* individuals spend considerable time looking around and sunbathing at the top of sand dunes (personal observation) to defend the field by showing threats and fighting to chase away intruders. In the face of threats from natural enemies, *P. mystaceus* can choose to run or bury themselves in the sand to avoid the enemy. Studies have shown that sand-burying behavior can reduce the risk associated with the occupation of exposed areas (Attum et al., 2007; Stellatelli et al, 2015). Therefore, the effect of fringes on the sand-diving performance of *P. mystaceus* has adaptive value. The repeated evolution of fringes among different groups of lizards and their relationship to specific environments (Luke, 1986; Halloy et al., 1998; Irschick and Jayne, 1998; Korff and McHenry, 2011) strongly support adaptive explanations involving movement (and burying) in sandy environments. In addition, toe fringe has been derived in the *Uma* genus (Etheridge and de Queiroz, 1986), which is probably due to adaptations to sand-bearing environments (Carothers, 1986). Molecular phylogenetic studies based on mitochondrial gene fragments have shown that *P. mystaceus* belongs to the primitive group in *Phrynocephalus* (Pang et al., 2003; Guo and Wang, 2007). Sand-diving behavior may be a primitive feature. Other *Phrynocephalus* species (with less developed fringes) can show sand-burying behaviors, but they have difficulty completing the task (Arnold, 1995). However, recent studies based on nuclear genes (nuDNA) suggest that the location of the base position of *P. mystaceus* is the result of interspecific hybridization and ancient mitochondrial gene infiltration. The common ancestor of *Phrynocephalus* probably preferred sandy substrates with the inclusion of clay or gravel. Climate change in the middle Miocene led to the migration of *Phrynocephalus* lizards into the desert, specifically the diffusion and adaptive evolution of large wind-blown dune habitats (Solovyeva et al., 2018). If this is the case, then the fringes of the *Phrynocephalus*, similar to those of *Uma*, are also the result of convergent evolution to adapt to the sand environment.

## Conclusion

The fringes of *P. mystaceus* play a significant role in its antipredation behavior in terms of maximum sprint speed. The presence of lateral fringes is more conducive to running in *P. mystaceus*. However, the species exhibits better sand-diving performance under the bilateral fringe state. Moreover, TFA can promote the burying performance of *P. mystaceus*. The fringes also play an important role in the selection of antipredation strategies. In the three fringe states, the predator avoidance strategy of *P. mystaceus* gradually shifted from sand diving to locomotor. It also shows that the fringes play a great role in sand-burying behavior.

## Animal Ethics

The following information was supplied relating to ethical approvals (i.e., approving body and any reference numbers): Specimens were collected following Guidelines for Use of Live Amphibians and Reptiles in Field Research (the Herpetological Animal Care and Use Committee (HACC) of the American Society of Ichthyologists and Herpetologists, 2004). This study was conducted in compliance with current laws on animal welfare and research in China and the regulations set by the Xinjiang Agricultural University. After the research was completed, the lizards were released where they were captured.

## Acknowledgments

We would like to thank the National Natural Science Foundation of China (31660613, 31260511) for support. Special appreciation goes to Xinqi WANG and Tong ZHOU from Xinjiang Agricultural University for their assistance in field work. We are also grateful to the reviewers for their useful and detailed advice.

## Author contributions

Conceptualization: P.Z., T.L., S.L. Methodology: P.Z., T.L., S.L. Formal analysis: P.Z. Investigation: P.Z., T.L., J.A., S.L. Writing - original draft: P.Z.

